# Cellsnp-lite: an efficient tool for genotyping single cells

**DOI:** 10.1101/2020.12.31.424913

**Authors:** Xianjie Huang, Yuanhua Huang

## Abstract

**Summary:** Single-cell sequencing is an increasingly used technology and has promising applications in basic research and clinical translations. However, genotyping methods developed for bulk sequencing data have not been well adapted for single-cell data, in terms of both computational parallelization and simplified user interface. Here we introduce a software, cellsnp-lite, implemented in C/C++ and based on well supported package htslib, for genotyping in single-cell sequencing data for both droplet and well based platforms. On various experimental data sets, it shows substantial improvement in computational speed and memory efficiency with retaining highly concordant results compared to existing methods. Cellsnp-lite therefore lightens the genetic analysis for increasingly large single-cell data.

**Availability:** The source code is freely available at https://github.com/single-cell-genetics/cellsnp-lite.

## 1 Introduction

Single-cell sequencing has become a powerful technology for disentangling heterogeneity in cell population at different levels, including genetics, transcriptome and epigenetics, hence has profound implications in basic research and clinical translations. The cellular genotype was primarily studied by single-cell DNA-seq (scDNA-seq) for detecting somatic mutations in tumors, clustering cells into clones and inferring their evolutionary dynamics (Navin, 2014). Recently, more evidence has been found that a subset of somatic mutations can also be observed in other single-cell probes, including scATAC-seq and full-length scRNA-seq (e.g., SMART-seq2), at both nuclear (McCarthy *et al*., 2020) and mitochondrial genomes (Ludwig *et al*., 2019). On the other hand, germline variants (a.k.a., single nucleotide polymorphisms, SNPs) are more widely observed in single-cell sequencing data, even in shallow droplet-based platforms, e.g., 10x Genomics, thanks to the large candidate list (around 7 million SNPs in human population with frequency > 5% (1000 Genomes Project Consortium, 2015)). Germline SNPs are not only perfect natural barcodes when multiplexing cells from multiple individuals (Huang *et al*., 2019), but also important in implying functional regulation via cellular eQTL analysis or allele specific expression (Cuomo *et al*., 2020), and allelic imbalance caused by copy number variation (Fan *et al*., 2018; Zaccaria and Raphael, 2020).

Genotyping methods for bulk sequencing sample are nearly mature with a decade of efforts and many methods remain effective when applying into single-cell sequencing data (Liu *et al*., 2019), including the successful BCFtools (Li *et al*., 2009; Li, 2011). However, there is lack of good adaption of these methods for single-cell data in terms of computational parallelization and simplified user interface. Here, we develop cellsnp-lite, a htslib (Li *et al*., 2009) based tool for genotyping in single cells. Htslib is a well developed, optimized and maintained package and is the core library used by BCFtools, hence we expect cellsnp-lite to give comparable accuracy as BCFtools in genotyping but higher efficiency and better convenience for single-cell data.

The goal of cellsnp-lite is to provide a user-friendly command line interface, with achieving high efficiency in both speed and memory. Therefore, it is designed as a light way allelic reads pileup with minimum filtering by keeping most data for customized downstream filtering and/or statistical modelling. We expect cellsnp-lite to be convenient in intermediate processing, e.g., for allelic ratio in copy number variations, and immediate useful for coarse analysis in less sensitive situation, e.g., for sample swap check.

## 2 Implementation

Cellsnp-lite is implemented in C/C++ and performs per cell genotyping, supporting both with (mode 1) and without (mode 2) given SNPs. In the latter case, heterozygous SNPs will be detected automatically. Cellsnp-lite is applicable for both droplet-based (e.g., 10x Genomics data) and well-based platforms (e.g., SMART-seq2 data). See **Table 1** for summary of these four options, and example alternatives in each mode.

**Table 1.** Cellsnp-lite genotype options and example alternatives

| Moe | SNPs | Bam files | Platform | Alternative |
| --- | --- | --- | --- | --- |
| Mode 1a | Given | pooled one | Droplet | VarTrix |
| Mode 1b | Given | each per cell | SMART-seq | BCFtools mpileup |
| Mode 2a | To detect | pooled one | Droplet | freebayes + VarTrix |
| Mode 2b | To detect | each per cell | SMART-seq | freebayes + mpileup |

Cellsnp-lite requires aligned reads as input, in bam / sam / cram file formats. Cell labels can be coded in the cell tag in a multiplexed bam file (droplet-based platforms) or specified by each per-cell bam file (well-based platforms). This flexibility also allows cellsnp-lite to work seamlessly on bulk samples, e.g., bulk RNA-Seq, by simply treating it as a well-based “cell”.

The pileup is performed per genome position, either for given SNPs (mode 1) or the whole chromosome (i.e., mode 2). All reads covering a query position will be fetched. By default, we discard those reads with low alignment quality, including MAPQ<20, aligned length<30nt, and FLAG with UNMAP, SECONDARY, QCFAIL (and DUP if UMI is not applicable). We then assign all these reads into each cell by hashmap for droplet-based sample (mode 1a or 2a) or direct assignment for well-based cells (mode 1b or 2b). Within each cell, we count the UMIs (if exists) or reads for all A, C, G, T, N bases. The REF and ATL alleles are taken from the input SNPs if given (i.e., in mode 1) otherwise by selecting the base with highest count as REF and second highest as ALT (mode 2).

When SNPs are given (mode 1), cellsnp-lite will perform parallel computing by splitting the input SNPs in-order and equally into multiple threads. Otherwise in mode 2, cellsnp-lite will compute in parallel by splitting the listed chromosomes, with each thread for one chromosome.

In all above scenarios, cellsnp-lite outputs sparse matrices for alternative allele, depths (i.e., REF and ALT alleles), and other alleles. If adding argument “--genotype”, cellsnp-lite will perform genotyping with the error model as presented in **Table 1** in (Jun *et al*., 2012), and output in VCF format with cell as samples.

## 3 Performance

### 3.1 High accuracy in pileup allelic counts

As discussed above, we aim for a light way pileup of allelic counts in large single-cell sequencing data. Hence, we mainly compared the pileup allelic counts between cellsnp-lite, cellsnp-Python, bcftools, vartrix (available at https://github.com/10XGenomics/vartrix; release version 1.1.16) and freebayes (Garrison and Marth, 2012). Unsurprisingly, we found cellsnp-lite gives identical allelic read counts compared to both its Python version and bcftools, as all use htslib for reads fetching (see settings in **Section 4.3** and **6.3** in **Supplementary file** for mode 1b and 2b on well-based cells, respectively).

In addition, for mode 1a with given SNPs on droplet-based data, we compared cellsnp-lite with vartrix, and found they are highly concordant (~97% SNPs with Pearson’s correlation coefficient >0.99, and >99.9% SNPs with mean absolute error < 0.01; **Supplementary Table S1-S4**). When there is no candidate SNPs given (mode 2), cellsnp-lite calls heterozygous SNPs directly. We found that in this setting, cellsnp-lite gives identical allelic counts compared to its mode 1 with given SNPs. Comparing to a commonly used alternative method, freebayes, for calling heterozygous SNPs from droplet-based scRNA-seq data, we found cellsnp-lite gives marginally higher accuracy (area under the precision recall curve, AUPRC: 0.964 vs 0.963; **Fig. S2**), where we treat the SNP arrays-based genotype as ground truth.

### 3.2 Substantial improvement on running speed

Thanks to well supported parallel computing, cellsnp-lite substantially outperforms existing methods in all above settings, with achieving around 6x to 14x speedups in droplet-based data (with large size: 10 to 100 GB per sample). When SNPs are given in mode 1a, cellsnp-lite is around 6x speedups and could save up to 90% peak memory compared to vartrix (**Fig. 1A** and **Fig. S1**). When lack of known SNPs in mode 2a, cellsnp-lite is about 7x ~ 14x speedups with no more than 2 times memory than freebayes in calling heterogynous SNPs (**Fig. 1C**).

**Figure 1.**
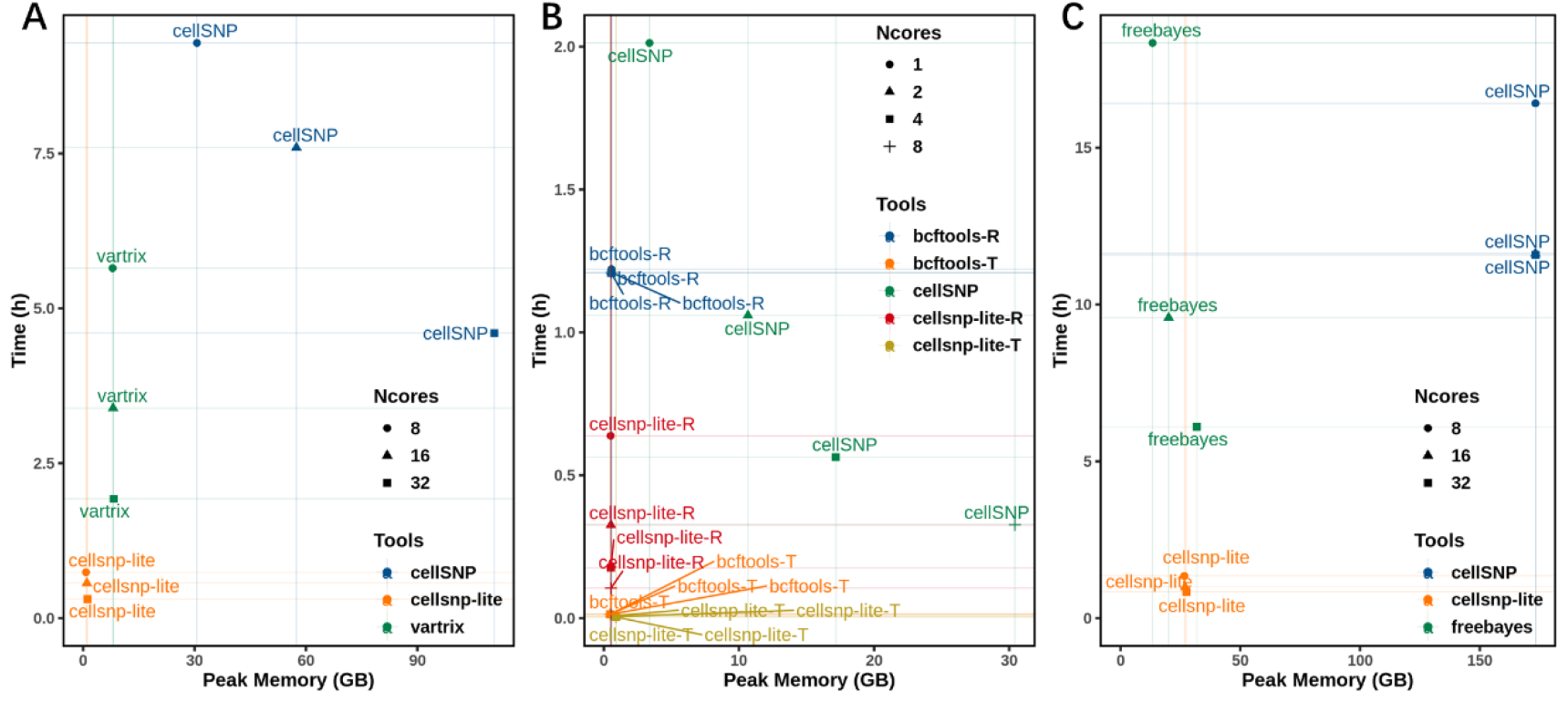
Cellsnp-lite showed high efficiency for genotyping in terms of running time and peak memory. CellSNP is the predecessor in Python of cellsnp-lite. **(A)** Comparison for mode 1a. Compared to vartrix, cellsnp-lite was about 6x speedups and could save up to 90% memory. Limited by huge memory usage, cellSNP could be slower than vartrix on a relatively big dataset. **(B)** Comparison for mode 1b. Cellsnp-lite could utilize multi-threading for processing query variants while bcftools mpileup could only use multi-threading for compression of the output stream. For either -*R* or -*T* option, cellsnp-lite was faster (about 2x ~ 12x speedups for -*R* and around 1.5 ~ 4x speedups for -*T*) than bcftools mpileup, even with single thread. Compared to bcftools mpileup, cellsnp-lite used slightly less memory with -*R* option while used no more than 2 times memory with -*T* option. For both cellsnp-lite and bcftools mpileup, the -*T* option was much (>30x) faster than -*R* option on this small dataset. Using large memory, cellSNP could be faster than bcftools mpileup -*R* option with many cores. **(C)** Comparison for mode 2a. Cellsnp-lite was the fastest among the three tools. Compared to freebayes, cellsnp-lite was about 7x ~ 14x speedups with no more than 2 times memory. The memory usage of freebayes gradually approached and eventually exceeded the one of cellsnp-lite as the number of threads increased. Limited by the huge memory usage, cellSNP gained little increase of speed with many threads.

Interestingly, cellsnp-lite also clearly outperforms bcftools mpileup on speed (with around 1.5x ~ 12x speedups) for well-based samples thanks to its better use of multi-threading as bcftools only uses multiple threads for writing file (**Fig. 1B** for given SNPs and **Fig. S3** for de-novo genotyping). We also noticed that since the bam files are small in well-based cells (50 to 500MB per cell), the bcftools *“-T”* option is more efficient to stream the whole bam file instead of to fetch each SNP through index file. In this particular setting with much less computing demand, cellsnp-lite (also with *T”* option) is still about 1.5x ~ 4x speedups than bcftools with using no more than 2 times memory (**Fig. 1B**).

## 4 Conclusion

Cellsnp-lite aims to pileup the expressed alleles in single-cell or bulk sequencing data with simple filtering. It has highly concordant results but substantially higher speed and less memory usage compared to other methods. Cellsnp-lite also provides simplified user interface and better convenience that supports parallel computing, cell barcode and UMI tags. Taken together, cellsnp-lite is expected to largely boost single-cell genetics analysis, especially considering the increasingly large size of single-cell data.

## Acknowledgements

We thank members in Oliver Stegle’s lab, especially Davis McCarthy and Hana Susak, for using and providing feedbacks on the predecessor CellSNP and current cellsnp-lite.

