## Supplementary data for "Cellsnp-lite: an efficient tool for genotyping single cells"

*To whom correspondence should be addressed.

### 1. Introduction

Cellsnp-lite, based on well supported package htslib (Li *et al.*, 2009), is designed for genotyping in single-cell sequencing data for both droplet and well based platforms. It has highly concordant results compared to other methods, but more time and memory efficient. Cellsnp-lite is open source and available on Github at https://github.com/single-cell-genetics/cellsnp-lite

### 2. Benchmarking Information

#### 2.1. Platform

**Server**

- PBS (Portable Batch System) v19.0.0 with 88 CPUs and 500G RAM;
- OS: CentOS 7.6.1810 (Core), Kernel: Linux 3.10.0-957.el7.x86_64

**Softwares**

All softwares below, except cellranger and vartrix which had standalone binaries, were packaged by and installed through conda v4.9.2.

- bcftools: v1.10.2 (using htslib v1.10.2)
- bgzip: v1.10.2 (using htslib v1.10.2)
- cellranger: v4.0.0
- cellSNP: v0.3.2
- cellsnp-lite: v1.2.0 (using htslib v1.10.2)
- freebayes: v1.3.2-dirty
- Python: v3.8.6
- R: v4.0.3
- samtools: v1.10.2 (using htslib v1.10.2)
- STAR: v2.7.6a
- vartrix: v1.1.16 (the binary zip file was downloaded from its Github repo from this link)

Cellsnp-lite is the C/C++ version of cellSNP. While some default settings have been changed in C/C++ version, the two tools should give identical results with the same settings.

#### 2.2. Scripts

- All benchmarking scripts are available on Github at https://github.com/single-cell-genetics/cellsnp-lite/tree/master/scripts/benchmark.
- A Python script memusg was used to compute running time and peak memory usage of a process.

#### 2.3. Datasets

**Demuxlet dataset**

The demuxlet dataset is a 10x scRNA-seq dataset which was published in demuxlet (Kang *et al.*, 2018).

- BAM file:
  demux.B.merged.bam (~19G).
- Barcode file:
  demux.B.barcodes.tsv contains 4246 cell barcodes.

**Souporcell dataset**

The souporcell dataset is a 10x scRNA-seq dataset which was published in souporcell (Heaton *et* *al.*, 2020).

- Fastq files:

  64 fastq files (~34G in total) of ENA sample SAMEA4810598 euts_1 were downloaded and then renamed by this script to follow the bcl2fastq file naming convention which was required by cellranger-count. The renamed fastq files were then taken as input of cellranger-count pipeline which aligned the fastq files to human genome reference hg19 (listed and described below) and outputted a BAM file and a barcode list file by this script.
- BAM file:

  soupc.bam (~31G) generated by cellranger-count pipeline mentioned above.
- Barcode file:

  soupc.barcodes.tsv generated by cellranger-count pipeline mentioned above.
- Genotyping array calls:

  HPSI0914i-euts_1.wec.gtarray.HumanCoreExome-12_v1_0.20160912.genotypes.vcf.gz (Gzipped VCF file, ~11M) contains genotype calls of euts_1 cell line that were generated by Infinium HumanExome BeadChip.

**Cardelino dataset**

The cardelino dataset is a Smart-seq2 scRNA-seq dataset which was published in cardelino (McCarthy *et al.*, 2020).

- Fastq files:

  The first 10 fastq files (~224M in total) of sample joxm were downloaded. The 10 files were generated by 5 runs ERR2806033 ~ ERR2806037 with paired fastq files for each run. The files were aligned to human genome reference hg19 (listed and described below) by star_aln.sh and then sorted by sam2bam.sh. The hg19 reference was indexed by star_index.sh beforehand.
- BAM file list:

  bam.lst contains absolute file paths to the 5 BAM files generated above.
- Sample list:

  sample.lst contains 5 sample names of the 5 BAM files generated above.
- Genotyping array calls:

  Genotype calls data of euts cell line, which could be used for further analysis, is available through the HipSci portal.

**Candidate SNP VCF file**

genome1K.phase3.SNP_AF5e2.chr1toX.hg19.noindel.nobiallele.nodup.vcf.gz (Gzipped VCF file, ~85M) contains ~7.36M SNPs.

It was converted from genome1K.phase3.SNP_AF5e2.chr1toX.hg19.vcf.gz, which was based on 1000 genome project and only kept SNPs on chr1-22 and X with minor allele frequency (MAF) > 0.05, by pre_filter_vcf.sh with command,

```
./pre_filter_vcf.sh -i genome1K.phase3.SNP_AF5e2.chr1toX.hg19.vcf.gz   \
    -o genome1K.phase3.SNP_AF5e2.chr1toX.hg19.noindel.nobiallele.nodup.vcf.gz
```

Then the SNPs of type indels, multi-alleles or duplicates were filtered by the command.

**Genome reference hg19**

The human genome reference hg19 file cellranger.hg19.3.0.0.fa was pre-compiled by 10x Genomics and can be download from this link.

#### 3. Benchmark of pileuping droplet-based datasets with given SNPs

##### 3.1. Softwares

Three tools were used for performance comparison for mode 1a:

- cellsnp-lite. (C/C++). Version v1.2.0 (using htslib v1.10.2)
- cellSNP. (Python). Version v0.3.2
- VarTrix. (Rust). Version v1.1.16

##### 3.2. Datasets

Previously described demuxlet dataset, souporcell dataset and candidate SNP VCF file.

##### 3.3. Tests

To evaluate the performance of cellsnp-lite on droplet-based datasets with given SNPs (mode 1a), two tests were performed to compare cellsnp-lite with vartrix on two 10x scRNA-seq datasets with the candidate SNP VCF file. Vartrix is an actively used software for extracting single cell variant information from 10x Genomics single cell data.

1. Pileup the demuxlet dataset with given SNPs
   script for running and script for comparison
2. Pileup the souporcell dataset with given SNPs
   script for running and script for comparison

*Evaluate performance of running time and peak memory*

Specifically, in both tests cellsnp-lite used *--cellTAG CB --UMItag UB --minCOUNT 1 --minMAF 0 --minLEN 0 --minMAPQ 20 --exclFLAG 772 --inclFLAG 0 --gzip* options, cellSNP used *--cellTAG CB --UMItag UB --minCOUNT 1 --minMAF 0 --minLEN 0 --minMAPQ 20 --maxFLAG 4096* options and vartrix used *--primary-alignments --umi --mapq 20 --bam-tag CB --scoring-method coverage --out-variants* options.

For comparison of running time and peak memory, all tests were repeated three times and average time and peak memory were used for plotting.

*Evaluate performance of accuracy*

As to the comparison of accuracy, file conversions were firstly performed on the outputs of the two tools. Cellsnp-lite outputted two matrices *AD.mtx* and *DP.mtx*. Vartrix also outputted two matrices *alt.mtx* and *ref.mtx*. Both the *AD.mtx* and *alt.mtx* contained depth information about alt allele of each variant for each cell. While *ref.mtx* stored depth information of ref allele, *DP.mtx* contained depth information of ref+alt alleles. This script was used for converting the *DP.mtx* to *REF.mtx* so that the new matrix has the same type of information as the one stored in *ref.mtx*. Then *AD.mtx* and *alt.mtx*, as well as *REF.mtx* and *ref.mtx*, were compared in pairs.

Two measurements that the Pearson correlation and Mean Absolute Error (MAE) were used for comparing the consistency of each pair of matrices separately.

#### 3.4. Results

##### Results of pileuping the demuxlet dataset with given SNPs

*Cellsnp-lite outperformed vartrix in running time and peak memory*

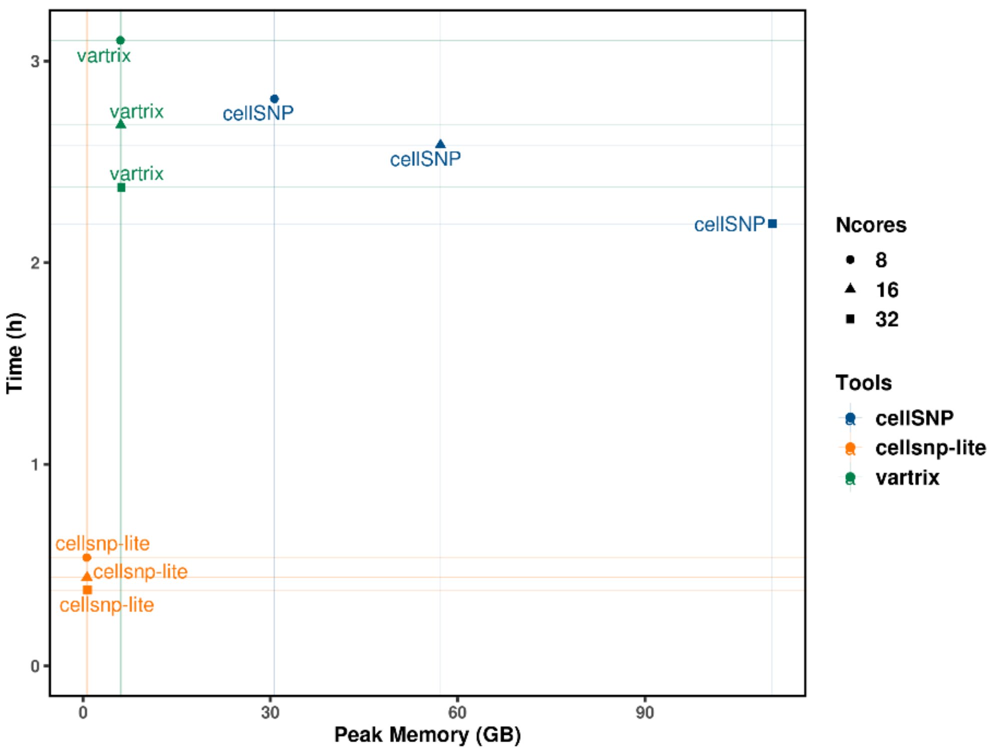

**Figure S1. Cellsnp-lite was the fastest and used the least memory among the three tools.** Compared to vartrix, cellsnp-lite was about 6x speedups and could save up to 90% peak memory. Although using much more memory, cellSNP was faster than vartrix on the demuxlet dataset which is a relatively small dataset.

*Cellsnp-lite had highly concordant results compared to vartrix*

**Table S1** and **Table S2** list separately the ratios of valid SNPs whose correlations and MAEs were above or below the thresholds for each matrix type. Both tables show high concordance between the pileup results of cellsnp-lite and vartrix on the demuxlet dataset.

**Table S1.** Pearson correlation between pileup results of cellsnp-lite and vartrix on demuxlet dataset.

| correlation (>) | matrix | valid SNP | total SNP | ratio (%) |
|---|---|---|---|---|
| 0.99 | Alt | 103752 | 105982 | 97.896 |
| 0.95 | Alt | 104452 | 105982 | 98.556 |
| 0.9 | Alt | 104829 | 105982 | 98.912 |
| 0.8 | Alt | 105340 | 105982 | 99.394 |
| 0.99 | Ref | 208306 | 213273 | 97.671 |
| 0.95 | Ref | 209858 | 213273 | 98.399 |
| 0.9 | Ref | 210808 | 213273 | 98.844 |
| 0.8 | Ref | 211941 | 213273 | 99.375 |

**Table S2.** MAE for pileup results of cellsnp-lite and vartrix on demuxlet dataset.

| MAE (<) | matrix | valid SNP | total SNP | ratio (%) |
|---|---|---|---|---|
| 0.01 | Alt | 105905 | 105982 | 99.927 |
| 0.05 | Alt | 105965 | 105982 | 99.984 |
| 0.1 | Alt | 105972 | 105982 | 99.991 |
| 0.2 | Alt | 105976 | 105982 | 99.994 |
| 0.01 | Ref | 213118 | 213273 | 99.927 |
| 0.05 | Ref | 213239 | 213273 | 99.984 |
| 0.1 | Ref | 213260 | 213273 | 99.994 |
| 0.2 | Ref | 213267 | 213273 | 99.997 |

**Results of pileuping the souporcell dataset with given SNPs**

*Cellsnp-lite outperformed vartrix in running time and peak memory*

See **Fig. 1A** in manuscript for the running time and peak memory used by cellsnp-lite and vartrix on the souporcell dataset.

*Cellsnp-lite had highly concordant results compared to vartrix*

**Table S3** and **Table S4** list separately the ratios of valid SNPs whose correlations and MAEs were above or below the thresholds for each matrix type. Both tables show high concordance between the pileup results of cellsnp-lite and vartrix on the souporcell dataset.

**Table S3.** Pearson correlation between pileup results of cellsnp-lite and vartrix on souporcell dataset.

| correlation (>) | matrix | valid SNP | total SNP | ratio (%) |
|---|---|---|---|---|
| 0.99 | Alt | 331182 | 343381 | 96.447 |
| 0.95 | Alt | 335744 | 343381 | 97.776 |
| 0.9 | Alt | 337903 | 343381 | 98.405 |
| 0.8 | Alt | 340694 | 343381 | 99.217 |
| 0.99 | Ref | 687808 | 719572 | 95.586 |
| 0.95 | Ref | 700403 | 719572 | 97.336 |
| 0.9 | Ref | 706228 | 719572 | 98.146 |
| 0.8 | Ref | 713296 | 719572 | 99.128 |

**Table S4.** MAE for pileup results of cellsnp-lite and vartrix on souporcell dataset.

| MAE (<) | matrix | valid SNP | total SNP | ratio (%) |
|---|---|---|---|---|
| 0.01 | Alt | 343116 | 343381 | 99.923 |
| 0.05 | Alt | 343322 | 343381 | 99.983 |
| 0.1 | Alt | 343353 | 343381 | 99.992 |
| 0.2 | Alt | 343366 | 343381 | 99.996 |
| 0.01 | Ref | 718893 | 719572 | 99.906 |
| 0.05 | Ref | 719442 | 719572 | 99.982 |
| 0.1 | Ref | 719510 | 719572 | 99.991 |
| 0.2 | Ref | 719555 | 719572 | 99.998 |

### 4. Benchmark of pileuping well-based dataset with given SNPs

#### 4.1. Softwares

Three tools were used for performance comparison for mode 1b:

- cellsnp-lite. (C/C++). Version v1.2.0 (using htslib v1.10.2)
- bcftools (mpileup). (C/C++). Version v1.10.2 (using htslib v1.10.2)
- cellSNP. (Python). Version v0.3.2

#### 4.2. Datasets

Previously described cardelino dataset and candidate SNP VCF file.

#### 4.3. Tests

To evaluate the performance of cellsnp-lite on well-based datasets with given SNPs (mode 1b), a test was performed to compare cellsnp-lite with bcftools mpileup on a smart-seq2 dataset with the candidate SNP VCF file. Bcftools mpileup is widely used for extracting variant information from bulk sequencing data and it could be applied to smart-seq data with proper settings.

1. Pileup the cardelino dataset with given SNPs
   script for running and script for comparison

*Evaluate performance of running time and peak memory*

Specifically, both cellsnp-lite and bcftools mpileup have two ways to specify locations, via *-R* or *-T* option. To access the next position, *-T* option streams the whole BAM while *-R* uses random access. For either *-R* or *-T* option, cellsnp-lite used *--cellTAG None --UMItag None --minCOUNT 1 --minMAF 0 --minLEN 0 --minMAPQ 20 --exclFLAG 1796 --inclFLAG 0 --gzip --genotype* options and bcftools mpileup used *-d 100000 -q 20 -Q 0 --incl-flags 0 --excl-flags 1796 -I -a AD,DP* options. CellSNP has only *-R* option and it used *--cellTAG None --UMItag None --minCOUNT 1 --minMAF 0 --minLEN 0 --minMAPQ 20 --maxFLAG 255* options.

For comparison of running time and peak memory, all tests were repeated three times and average time and peak memory were used for plotting.

*Evaluate performance of accuracy*

As to the comparison of accuracy, file conversions were firstly performed on the outputs of the two tools. Cellsnp-lite outputted two matrices *AD.mtx* and *DP.mtx* and this script was used for converting the *DP.mtx* to *REF.mtx*. Bcftools mpileup outputted one VCF file and this script was used for extracting the read depth of ref and alt alleles from the VCF to generate the *ref.mtx* and *alt.mtx* separately. Then *AD.mtx* and *alt.mtx*, as well as *REF.mtx* and *ref.mtx*, were compared in pairs.

Two measurements that the Pearson correlation and Mean Absolute Error (MAE) were used for comparing the consistency of each pair of matrices separately.

#### 4.4. Results

**Results of pileuping the cardelino dataset with given SNPs**

*Cellsnp-lite was faster than bcftools mpileup*

See **Fig. 1B** in manuscript for the running time and peak memory used by cellsnp-lite and bcftools mpileup on the cardelino dataset.

*Pileup results of cellsnp-lite were totally the same with bcftools mpileup*

The two pairs of matrices that *REF.mtx* & *ref.mtx* and *AD.mtx* & *alt.mtx* were compared separately with Linux shell command ***diff*** and it showed that each pair of matrices were completely the same.

### 5. Benchmark of pileuping droplet-based dataset without given SNPs

#### 5.1. Softwares

Three tools were used for performance comparison for mode 2a:

- cellsnp-lite. (C/C++). Version v1.2.0 (using htslib v1.10.2)
- cellSNP. (Python). Version v0.3.2
- freebayes. (C/C++). Version v1.3.2-dirty

#### 5.2. Datasets

Previously described souporcell dataset.

#### 5.3. Tests

Cellsnp-lite uses a simple model (Jun *et al.*, 2012; Kang *et al.*, 2018) and filtering strategy for identifying candidate SNPs. To evaluate the performance of cellsnp-lite on droplet-based datasets without given SNPs (mode 2a), a test was performed to compare cellsnp-lite with freebayes on a 10x scRNA-seq dataset. Freebayes is widely used for genotyping on bulk sequencing dataset and it could be applied to single cell sequencing data with proper settings. The combination of freebayes and vartrix is an alternative to cellsnp-lite mode 2a.

1. Call candidate SNPs from the souporcell dataset
   script for running and script for comparison
2. Compare pileup results of mode 2a with mode 1a on the souporcell dataset
   script for running and script for comparison

**Call candidate SNPs from the souporcell dataset**

*Evaluate performance of running time and peak memory*

Specifically, the first test compared the performance of cellsnp-lite with freebayes on the souporcell dataset. The three tools called SNPs from the souporcell dataset. Cellsnp-lite used *--chrom 1,2,3,4,5,6,7,8,9,10,11,12,13,14,15,16,17,18,19,20,21,22,X,Y --cellTAG None --UMItag None --minCOUNT 1 --minMAF 0 --minLEN 0 --minMAPQ 20 --exclFLAG 772 --inclFLAG 0 --gzip --genotype* options. To speed up, the parallele version of freebayes, named freebayes-parallel, was used according to its recommendation on Github. Freebayes-parallel used the command *freebayes-parallele <(fasta_generate_regions.py <fa.fai> 100000)* with *--genotype-qualities --use-duplicate-reads* options. CellSNP used *--cellTAG None --UMItag None --minCOUNT 1 --minMAF 0 --minLEN 0 --chrom 1,2,3,4,5,6,7,8,9,10,11,12,13,14,15,16,17,18,19,20,21,22,X,Y --minMAPQ 20 --maxFLAG 4096* options.

For comparison of running time and peak memory, all tests were repeated three times and average time and peak memory were used for plotting.

*Evaluate performance of accuracy*

The euts_1 cell line, which was sequenced and used by souporcell dataset, had genotype calls that were generated from genotyping array. The genotype calls were used as ground truth. Only the shared SNPs of tool(s) and ground truth were used for downstream analysis. Besides, cellsnp-lite would not know exactly the real ref alleles without given SNPs so the TGT (Translated Genotype, e.g., A/A) was used for comparing instead of GT (Genotype, e.g., 0/0). Precision-Recall curves were plotted using GQ (Genotype Quality) as scores.

**Compare pileup results of mode 2a with mode 1a on the souporcell dataset**

The second test compared the pileup results of cellsnp-lite mode 2a with mode 1a on the souporcell dataset. Cellsnp-lite mode 2a used *--cellTAG CB --UMItag UB --minCOUNT 1 --minMAF 0 --chrom 1,2,3,4,5,6,7,8,9,10,11,12,13,14,15,16,17,18,19,20,21,22,X,Y --minLEN 0 --minMAPQ 20 --exclFLAG 772 --inclFLAG 0 --gzip --genotype* options. The options used by cellsnp-lite mode 1a were described previously. Containing all SNPs extracted from the input BAM file, the output file of cellsnp-lite mode 2a was firstly intersected with the given SNPs used in mode 1a. The shared SNPs of the two modes were compared.

#### 5.4. Results

##### Results of calling candidate SNPs on the souporcell dataset

*Cellsnp-lite was faster than freebayes*

See **Fig. 1C** in manuscript for the running time and peak memory used by cellsnp-lite and freebayes calling on the souporcell dataset.

*Cellsnp-lite had highly concordant results compared to freebayes on calling heterozygous SNPs*

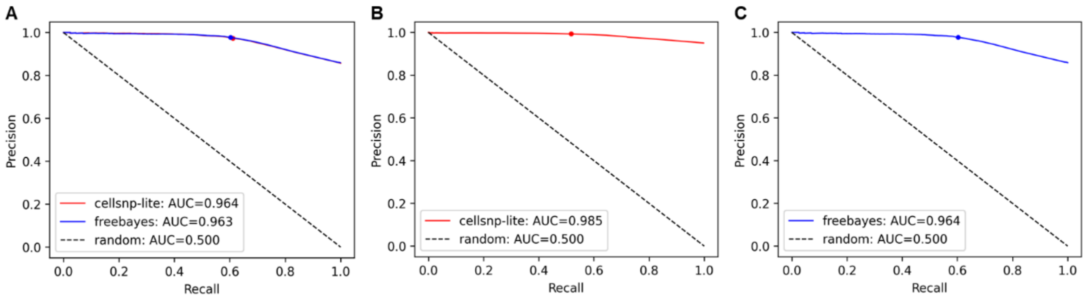

**Figure S2. Cellsnp-lite was highly consistent with freebayes on calling heterozygous SNPs.** The red or blue dot denoted the result when GQ equals 20 for cellsnp-lite and freebayes separately. **(A)** Results for the 11976 shared SNPs (among them 5737 are homozygous) of cellsnp-lite, freebayes and ground truth. The curves and GQ20 dots of the two tools almost coincided with each other and both had high AUC. **(B)** Results for the 160032 shared SNPs of cellsnp-lite and ground truth. Cellsnp-lite showed a bit higher AUC compared to (A) as 147363 homozygous SNPs existed in the shared part. **(C)** Results for the 11995 shared SNPs (among them 5754 are homozygous) of freebayes and ground truth. The AUC was the same with (A) as the SNPs in (A) and (C) were almost the same.

##### Results of comparing pileup results of mode 2a with mode 1a on the souporcell dataset

The *AD.mtx* and *DP.mtx* outputted by cellsnp-lite mode 2a were firstly intersected with the given SNPs used in mode 1a and then were compared with the corresponding matrices of mode 1a by running Linux shell command ***diff***. It showed that each pair of matrices were completely the same.

### 6. Benchmark of pileuping well-based dataset without given SNPs

#### 6.1. Softwares

Three tools were used for performance comparison for mode 2b:

- cellsnp-lite. (C/C++). Version v1.2.0 (using htslib v1.10.2)
- bcftools (mpileup). (C/C++). Version v1.10.2 (using htslib v1.10.2)
- cellSNP. (Python). Version v0.3.2

#### 6.2. Datasets

Previously described cardelino dataset.

#### 6.3. Tests

To evaluate the performance of cellsnp-lite on well-based datasets without given SNPs (mode 2b), a test was performed to compare cellsnp-lite with bcftools mpileup on a smart-seq2 dataset. Bcftools mpileup is widely used for extracting variant information from bulk sequencing data and it could be applied to smart-seq data with proper settings. The combination of freebayes and bcftools mpileup is an alternative to cellsnp-lite mode 2b.

1. Pileup the cardelino dataset without given SNPs
   script for running and scripts 1 & 2 for comparison

*Evaluate performance of run time and peak memory*

Specifically, cellsnp-lite used *--cellTAG None --UMItag None --minCOUNT 1 --minMAF 0 --minLEN 0 --chrom 1,2,3,4,5,6,7,8,9,10,11,12,13,14,15,16,17,18,19,20,21,22,X,Y --minMAPQ 20 --exclFLAG 1796 --inclFLAG 0 --gzip --genotype* options, bcftools mpileup used *-d 100000 -q 20 -Q 0 --incl-flags 0 --excl-flags 1796 -I -a AD,DP* options and cellSNP used *--cellTAG None --UMItag None --minCOUNT 1 --minMAF 0 --minLEN 0 --minMAPQ 20 --maxFLAG 255 --chrom 1,2,3,4,5,6,7,8,9,10,11,12,13,14,15,16,17,18,19,20,21,22,X,Y* options.

For comparison of running time and peak memory, all tests were repeated three times and average time and peak memory were used for plotting.

*Evaluate performance of accuracy*

Two steps were taken to evaluate the consistency between cellsnp-lite with bcftools mpileup for mode 2b and the consistency between cellsnp-lite mode 2b with mode 1b.

Firstly, the read depths for each base of {A, C, G, T} were extracted from the outputted VCF to generate two SNP x Cell matrices for cellsnp-lite and bcftools mpileup separately. Note that base N was not included in comparison as bcftools mpileup did not count N while cellsnp-lite did. Secondly, the output file of cellsnp-lite mode 2b was intersected with the given SNPs used in mode 1b and the shared SNPs of the two modes were compared.

#### 6.4. Results

##### Results of pileuping the cardelino dataset without given SNPs

*Cellsnp-lite outperformed bcftools mpileup in running time and peak memory*

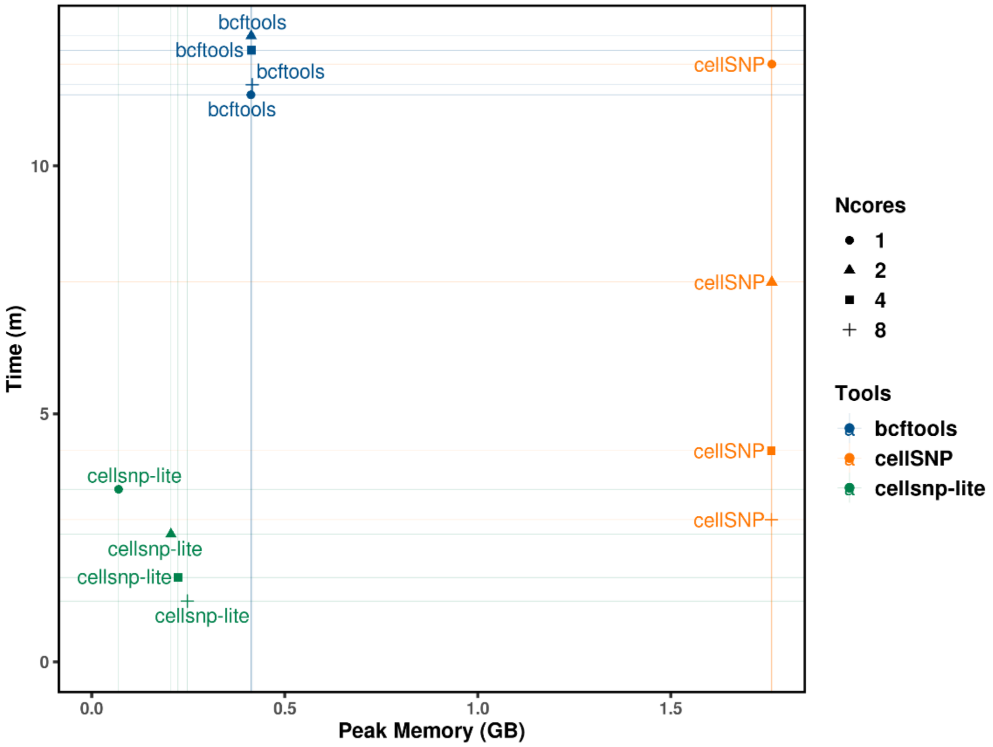

**Figure S3. Cellsnp-lite was the fastest and used the least memory among the three tools.** Cellsnp-lite could utilize multi-threading for processing query variants while bcftools mpileup could only use multi-threading for compression of the output stream. Using less memory, cellsnp-lite was around 3x ~ 9x speedups than bcftools mpileup, even with single thread. CellSNP could be faster than bcftools mpileup with many cores and large memory.

*Pileup results of cellsnp-lite were totally the same with bcftools mpileup*

The two SNP x Cell matrices of read depths for each base {A, C, G, T} were compared directly with Linux shell command ***diff*** and it showed that they were completely the same.

*Pileup results for the shared SNPs of cellsnp-lite mode 2b and mode 1b were totally the same*

The *AD.mtx* and *DP.mtx* outputted by cellsnp-lite mode 2b were firstly intersected with the given SNPs used in mode 1b and then were compared with the corresponding matrices of mode 1b by running Linux shell command ***diff***. It showed that each pair of matrices were completely the same.
